# Haplotype graph analysis of *PdR1* uncovers resistance diversity to Pierce’s Disease in *Vitis arizonica* and its hybrids

**DOI:** 10.64898/2025.12.23.696282

**Authors:** Mélanie Massonnet, Mirella Zaccheo, Noé Cochetel, Rosa Figueroa-Balderas, Summaira Riaz, Dario Cantu

## Abstract

Previous genetic mapping studies indicate that multiple haplotypes of the *Pierce’s disease (PD) Resistance 1 (PdR1)* locus occur in *Vitis arizonica* and its hybrids. To characterize sequence diversity at this locus, we assembled chromosome-scale diploid genomes for four PD-resistant (PD-R) accessions: b43-17 (*PdR1a⁺/PdR1b⁺*), the backcross 07744-094 (*PdR1c⁺/PdR1⁻*), b46-43 (*PdR1e⁺/PdR1f⁺*), and b42-26 (*PdR1⁻/PdR1⁻*), which displays quantitative PD resistance not associated with *PdR1*. Haplotype resolution of *PdR1a*, *PdR1b*, *PdR1c*, and *PdR1e* revealed substantial variation in intergenic repeat content and gene composition between *PdR1* and their alternative haplotype at the *PdR1* locus not associated with PD resistance phenotype (*PdR1*^-^), as well as among *PdR1* haplotypes, demonstrating extensive sequence diversity at the *PdR1* locus. Sequence graph analysis uncovered substantial structural divergence concentrated in approximately one quarter of the locus, together with smaller-scale variation across haplotypes. This analysis identified *PdR1*-specific graph nodes, showing that *PdR1a* and *PdR1b* share most of their *PdR1*-specific features, whereas *PdR1c* contains the highest number of private nodes, followed by *PdR1e*. Integration of sequence graph features with gene expression data further refined a set of defense-related candidate genes within *PdR1c*. Together, these results identify candidate genes for functional validation and indicate that multiple resistance determinants co-localized within the *PdR1* locus may contribute to PD resistance, highlighting opportunities for targeted genetic improvement strategies.

## Introduction

Pierce’s disease (PD) is caused by the bacterium *Xylella fastidiosa* ssp. *fastidiosa* (hereafter *Xf*), which colonizes grapevine xylem. In California, PD remained limited until the late 1990s when the introduction of the non-native glassy-winged sharpshooter (*Homalodisca vitripennis*) in the southern part of the state led to a rapid rise in disease incidence (Rapicavoli et al., 2018). These insects, along with the native blue-green sharpshooter (*Graphocephala atropunctata*) and **s**pittlebug species (family Cercopidae), are principal vectors of *Xf.* All *Vitis vinifera* cultivars are susceptible to PD, whereas some wild grape species from the southern US and northern Mexico, including *V. arizonica* and its hybrids, are PD-resistant.

In grapes, *Xf* colonization triggers the formation of balloon-shaped tyloses in the xylem, which help block pathogen spread and limit air embolisms (Castro et al., 2021). While this defense response is effective in PD-resistant grapes, tylose formation is spatially and temporally misaligned with *Xf* colonization in PD-susceptible cultivars, leading to extensive vessel occlusion (Sun et al., 2013). Coupled with systemic *Xf* spread through bacterial biofilms, excess tylose development sharply reduces stem hydraulic conductivity in susceptible grapevines (Rapicavoli et al., 2018). This results in leaf scorching, berry desiccation, irregular periderm development, abnormal leaf abscission, and vine death (Rapicavoli et al., 2018).

PD management mostly relies on the use of insecticides to control vectors and the removal of infected plants (Haviland et al., 2021). Despite these practices, annual production loss is estimated at $56.1 million per year in California (Tumber et al., 2014). In addition, growing application restrictions and emerging pesticide resistance have made this approach unsustainable (Haviland et al., 2021). Therefore, the long-term solution is the development of grape cultivars with strong and durable PD resistance combined with desirable agronomic and enological traits. Such resistance can be achieved through the introgression of functionally diverse PD resistance-associated genes or alleles into *V. vinifera* (Michelmore et al., 2013).

One resistance (*R*) locus located on chromosome 14, *PD Resistance 1* (*PdR1*), was identified in grapes through genetic mapping (Krivanek et al., 2006; Riaz et al., 2006). Using marker-based maps and tightly linked molecular markers, putative *PdR1* haplotypes were distinguished among *V. arizonica* accessions and their hybrids based on marker size polymorphisms rather than sequence data (Riaz et al., 2008; Riaz et al., 2018). The presence of multiple marker-defined haplotypes at the same locus (*PdR1a-f*) suggests that PD resistance may involve different genes or allelic variants within *PdR1* (**Table 1**). The diploid genome of *V. arizonica* b40-14, which carries *PdR1c* and *d* (*PdR1c*^+^*/PdR1d*^+^), together with a genome-wide association and gene expression profiling, enabled the identification of four candidate *R* genes encoding extracellular receptors: two leucine-rich repeat (LRR) receptor-like proteins (RLPs), one LRR-kinase (LRR-RLK), and one lysin motif (LysM)-RLK (Morales-Cruz et al., 2023). However, limited haplotype information prevented confirmation of the locus phasing and whether the studied *PdR1* haplotype corresponded to the *c* or *d* haplotype of b40-14.

**Table 1:**
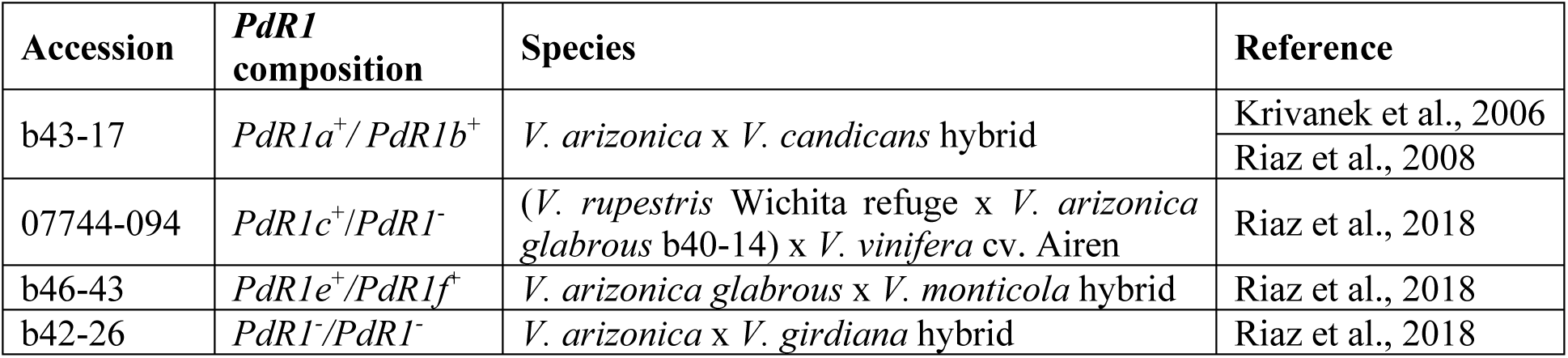
List of the PD-resistant grape accessions whose genome was sequenced in this study.

*PdR1b* has already been deployed in the University of California Davis grape breeding program and underlies the five PD-resistant cultivars released in 2019 (https://fps.ucdavis.edu/newsarticle.cfm?newsid=57). These cultivars have generated significant industry interest, motivating efforts to characterize additional *PdR1* haplotypes that could further strengthen resistance and improve its durability in future breeding. In this study, to better understand the range of resistance encoded at this locus, we assess sequence diversity among five *PdR1* haplotypes and its impact on genes potentially involved in *Xf* resistance. We sequenced, assembled, and annotated diploid chromosome-scale genomes for four PD-resistant accessions: b43-17 (*PdR1a*^+^*/ PdR1b*^+^), the backcross 07744-094 (*PdR1c*^+^/*PdR1*^-^), b46-43 (*PdR1e*^+^*/PdR1f*^+^), and b42-26 (*PdR1*^-^/*PdR1*^-^) which displays quantitative PD resistance but not associated with *PdR1* (**Table 1**) (Riaz et al., 2018). After localizing the *PdR1* haplotypes using marker data and phasing them, we compared the gene content potentially associated with *Xf* resistance across five *PdR1* haplotypes and three alternative haplotypes. *PdR1*^-^ will be used to describe the alternative haplotype at the *PdR1* locus not associated with PD-resistance phenotype. A sequence graph built from both *PdR1* and *PdR1*^-^ haplotypes allowed us to identify variants specific to the *PdR1* haplotypes, compare these variants across haplotypes, and determine which ones altered the coding sequences of defense-related genes. Finally, we show how the information of the sequence graph, coupled with gene expression, can help to narrow down candidate genes among the refined region of *PdR1c* (Morales-Cruz et al., 2023).

## Methods

### Plant material

Young leaves from the accessions b43-17 (*PdR1a*^+^*/ PdR1b*^+^), 07744-094 (*PdR1c*^+^/*PdR1*^-^), b46-43 (*PdR1e*^+^*/PdR1f*^+^), and b42-26 (*PdR1*^-^/*PdR1*^-^) were collected, and immediately ground in liquid nitrogen.

### DNA extraction, library preparation, and sequencing

High-molecular weight genomic DNA was isolated from the young leaves of the four accessions like in Chin *et al*. (Chin et al., 2016). DNA concentration was evaluated using the DNA High Sensitivity Kit on a Qubit 2.0 Fluorometer (Life Technologies, CA, USA), and DNA purity with a Nanodrop 2000 spectrophotometer (Thermo Scientific, IL, USA). Assessment of the DNA integrity was done using FEMTO Pulse system (Agilent, CA, USA). Preparation of the HiFi libraries was performed using the SMRTbell Express Template Prep Kit 2.0 (Pacific Biosciences, CA, USA) following the manufacturer’s protocol. Libraries were size-selected using 3.1 X v/v of 35% Ampure PB beads (Pacific Biosciences, CA, USA), and sequenced on a PacBio Sequel II platform (DNA Technology Core Facility, University of California, Davis, CA, USA). Sequencing resulted in 54, 54, 57, and 40X of coverage of a 500-Mbp haplotype for b43-17 (*PdR1a*^+^*/ PdR1b*^+^), 07744-094 (*PdR1c*^+^/*PdR1*^-^), b46-43 (*PdR1e*^+^*/PdR1f*^+^), and b42-26 (*PdR1*^-^/*PdR1*^-^), respectively (**Table S1**).

### Genome assembly

HiFi reads were assembled using hifiasm (Cheng et al. 2021) with the option “-n 13” for b43-17 (*PdR1a*^+^*/ PdR1b*^+^), 07744-094 (*PdR1c*^+^/*PdR1*^-^), and b42-26 (*PdR1*^-^/*PdR1*^-^), and “-n 25” for b46-43 (*PdR1e*^+^*/PdR1f*^+^) (**Table S2**). Diploid chromosome-scale pseudomolecules were scaffolded from contigs using HaploSync v1.0 (Minio et al., 2022) and consensus grape synteny map (Cochetel et al., 2025) (**Table S3**). Telomere repeat units were searched as described in Massonnet *et al*. (Massonnet et al., 2025) using TIDK v.0.2.63 (https://github.com/tolkit/telomeric-identifier). Gene space completeness of the genome assemblies was evaluated using BUSCO v.5.4.7 with the library embryophyte_odb10 (Waterhouse et al., 2018).

### Genome annotation

Repeat and gene annotations were performed as in Massonnet *et al*. (Massonnet et al., 2025). Repeat annotation was made using RepeatMasker v.4.1.7-p1 (Smit et al., 2013) and a grape repeat library (Minio et al., 2019). PN40024 V5.1 predicted protein-coding gene sequences (https://grapedia.org/t2t_annotation/), along with the high-quality transcripts obtained from the Iso-Seq full-length reads of *V. arizonica* b40-14 (Cochetel et al., 2023) and *V. vinifera* cv. Cabernet Sauvignon (Minio et al., 2019), were used as transcriptomic evidence for the gene annotation. For each accession, a set of high-quality gene models was created using the PASApipeline v2.5.3 (https://github.com/PASApipeline/PASApipeline), GMAP v.2024-11-20 (Wu and Watanabe, 2005), pblat v.2.5.1 (Wang and Kong, 2019) and the transcriptomic evidence aforementioned. The set of high-quality gene models was then used as input for the *ab initio* predictors Augustus v.3.5.0 (Stanke et al., 2006) and GeneMark v.3.68_lic (Lomsadze et al., 2005). Consensus models from the gene models produced by the *ab initio* predictions and PASA were made using EvidenceModeler v.2.1.0 (Haas et al., 2008). Gene models encoding protein sequences with a length less than 50 amino acids, or in-frame stop codons, or lacking a start methionine or a stop codon, were removed.

### Functional annotation

Predicted proteins of the four genomes were aligned onto the proteins of *Arabidopsis thaliana* (Araport11_genes.201606.pep.fasta; https://www.arabidopsis.org/download/index.jsp) using BLASTP v.2.12.0+ (Camacho et al., 2009). Homologous proteins were identified by: first, keeping the alignments with an identity greater than 30%, and both reference:query and query:reference coverages between 75% and 125%; second, selecting the alignment with the highest product of identity by query coverage by reference coverage for each grape protein. Same method was used to determine homologous proteins in *V. vinifera* PN40024 genome V1 annotation and V5.1 annotations (Jaillon et al., 2007; Shi et al., 2023).

To identify protein domains, hmmsearch from HMMER v.3.3.1 (Eddy, 2011) was used with the Pfam-A Hidden Markov Models (HMM) database (El-Gebali et al., 2019) (downloaded on March 12^th^, 2025). Protein domains with an independent E-value less than 1.0, and at least 50% of the HMM, were selected (**Tables S4-S7**).

Transmembrane helices were predicted with TMHMM2 v2.0c (Krogh et al., 2001).

### *PdR1* region localization

The five *PdR1* haplotypes and the three *PdR1*^-^ haplotypes were located by *in silico* amplification of two *PdR1*-associated markers, VMCNg2b7.2 and UDV095 (Riaz et al., 2008), on the genome assemblies using dipr (https://github.com/douglasgscofield/dispr).

### Haplotyping

To identify *PdR1c* in 07744-094 (*PdR1c*^+^/*PdR1*^-^) and *PdR1b* in b43-17 (*PdR1a*^+^*/ PdR1b*^+^), we used short DNA-seq reads from *V. arizonica* b40-14 (*PdR1c*^+^/*PdR1d*^+^) and five backcrossed cultivars carrying *PdR1b*: Ambulo blanc, Caminante blanc, Camminare noir, Errante noir and Paseante noir (Morales-Cruz et al., 2023), and a coverage analysis of unambiguous alignments. Adapter sequence removal and quality read selection was performed using Trimmomatic v.0.36 (Bolger et al. 2014) with the following parameters: “LEADING:3 TRAILING:3 SLIDINGWINDOW:10:20 MINLEN:36 CROP:150”. High-quality reads of *V. arizonica* b40-14 and the five backcrossed cultivars were aligned on 07744-094 and b43-17 genomes, respectively, using BWA v0.7.17 (Li and Durbin, 2009) with the parameter “-a” to report all multiple alignments. Unambiguous alignments (i.e., with no mismatches) were retained with bamtools filter v.2.5.1 (Barnett et al., 2011) and the option “NM:0”. Base coverage was obtained using genomecov from BEDTools v2.29.1 (Quinlan, 2014). Coverage in repetitive elements was removed using BEDTools intersect. Median coverage per window of 10-kbp was computed by BEDTools map. Median coverage per 10-kbp window was normalized by dividing by the sequencing coverage of the accessions. SNPs and INDELs between the two haplotypes of b43-17 were called using NUCmer from MUMmer v.4.0.0 (Marçais et al., 2018) with the “--mum” option and show-snps with the parameters “-Clr”.

### Sequence graph construction

Sequence graph of the five *PdR1* haplotypes and the three *PdR1*^-^ alternative haplotypes were built using the nf-core/pangenome pipeline (Heumos et al., 2024) with NextFlow v.25.04.6 (Di Tommaso et al., 2017) and the parameters “-r 1.1.2-profile singularity”. 2D representation was generated using ODGI v0.9.3 (Guarracino et al., 2022).

### Alignment of PD resistance-associated kmers

PD resistance-associated kmers from *V. arizonica* were retrieved from Morales-Cruz *et al*. (Morales-Cruz et al., 2023). They were mapped on the genome of 07744-094 (*PdR1c*^+^/*PdR1*^-^) using BLASTN v.2.12.0+ (Camacho et al., 2009) and a word size of 8 bp. Full-length and perfect alignments (i.e. 100% coverage and identity) were retained.

### Gene expression analysis

RNA-seq reads of three PD-resistant 07744 lines were retrieved from NCBI BioProject PRJNA956994 (Morales-Cruz et al., 2023). High-quality reads were obtained using Trimmomatic v.0.36 (Bolger et al., 2014) with the following parameters: “LEADING:3 TRAILING:3 SLIDINGWINDOW:10:20 MINLEN:36” (**Table S11**). Transcript abundance was assessed with Salmon v.1.5.1 (Patro et al., 2017). A transcriptome index file was built using the protein-coding sequences of 07744-094 and its genome as decoy, using the parameters “-k 31 --keepDuplicates”. Transcript abundance was quantified with the parameters “--gcBias --seqBias --validateMappings”. Gene expression in transcript per million (TPM) was extracted from quantification files with the R package tximport v.1.30.0 (Soneson et al., 2016). Differential gene expression analysis was performed using DESeq2 v.1.42.1 (Love et al., 2014).

## Results

### Diploid chromosome-scale genome assemblies

To characterize the sequence diversity among the *PdR1* haplotypes, we sequenced the genomes of three PD-resistant *V. arizonica* hybrids: *V. arizonica* x *V. candicans* hybrid b43-17 (*PdR1a*^+^*/ PdR1b*^+^), *V. arizonica glabrous* x *V. monticola* hybrid b46-43 (*PdR1e*^+^*/PdR1f*^+^), and *V. arizonica* x *V. girdiana* hybrid b42-26 (*PdR1*^-^/*PdR1*^-^) which has quantitative PD resistance not associated with *PdR1* (**Table 1**) (Riaz et al., 2018), using long-read DNA sequencing (PacBio HiFi reads; **Table S1**). In addition, we sequenced the genome of the backcross line 07744-094 to obtain the *PdR1c* haplotype from *V arizonica* b40-14 (*PdR1c*^+^/*PdR1d*^+^), because in the available b40-14 genome the two *PdR1* haplotypes are fragmented and not haplotype-phased (**Figure S1**) (Morales-Cruz et al., 2023).

Draft diploid genomes were composed of an average of 192 ± 43.4 contigs (**Table S2**). After pseudomolecule reconstruction, each set of 19 chromosomes spanned 491.9 ± 67.8 Mbp, with unanchored contigs accounting for only 1.7 ± 0.8% of the assembly (16.7 ± 8.1 Mbp) (**Table 2**; **Table S3**). On average, 99.2 ± 0.2% of expected single-copy orthologs (BUSCOs) were detected in each set of 19 chromosomes, indicating a high completeness. Furthermore, we detected telomeric repeats at both end of 24.5 ± 6.2 chromosomes and at one end of 12.3 ± 5.8 chromosomes out of 38. These results indicate that the four haplotype-resolved assemblies are highly contiguous and complete.

**Table 2:** Statistics of the four genome assemblies. Abbreviations: Hap, Haplotype; Unp, Unplaced contigs.

| Accession | b43-17 |  |  | 07744-094 |  |  | b46-43 |  |  | b42-26 |  |  |
| --- | --- | --- | --- | --- | --- | --- | --- | --- | --- | --- | --- | --- |
| *PdR1* composition | *PdR1a<sup>+</sup>/PdR1b<sup>+</sup>* |  |  | *PdR1c<sup>+</sup>/PdR1<sup>-</sup>* |  |  | *PdR1e<sup>+</sup>/PdR1f<sup>+</sup>* |  |  | *PdR1<sup>-</sup>/PdR1<sup>-</sup>* |  |  |
| Haplotype | Hap1 | Hap2 | Unp | Hap1 | Hap2 | Unp | Hap1 | Hap2 | Unp | Hap1 | Hap2 | Unp |
| Assembly length (Mbp) | 502.1 | 488.3 | 15.4 | 484.6 | 491.8 | 13.7 | 501.1 | 489.8 | 9.4 | 493.6 | 484.1 | 28.1 |
| Number of sequences | 19 | 19 | 81 | 19 | 19 | 70 | 19 | 19 | 56 | 19 | 19 | 93 |
| Telomeres | 29 | 24 | 0 | 33 | 35 | 0 | 29 | 30 | 0 | 33 | 33 | 0 |
| Number of genes (x1,000) | 39.2 | 38.3 | 1.0 | 37.5 | 36.1 | 3.3 | 32.1 | 31.4 | 0.8 | 33.1 | 32.4 | 2.1 |
| Repeats (%) | 56.3 | 56.0 | 79.9 | 56.5 | 56.0 | 76.8 | 57.3 | 56.6 | 69.7 | 56.1 | 55.8 | 78.9 |
| BUSCOs (%) | 99.2 | 98.9 | 1.1 | 99.4 | 99.5 | 0.4 | 99.1 | 99.1 | 0.1 | 99.1 | 99.1 | 1.8 |

### *PdR1* localization and haplotyping

The boundaries of *PdR1* and the *PdR1*^-^ haplotypes were located through *in silico* amplification of the *PdR1*-linked markers VMCNg2b7.2 and UDV095 (Riaz et al., 2008) (**Table S8**). Three additional markers (Ch14-70, Ch14-77, and Ch14-02) were used to confirm their localization (Riaz et al., 2018). Amplification revealed amplicon-size variation between accessions for all markers. Variation within accessions, reflecting differences between the two homologous chromosomes, was also observed for at least one marker in each accession, except b46-43, whose two haplotypes produced identical amplicon sizes for all markers (**Table S8**). The size of the *PdR1* region varied across the four grape accessions, providing further evidence of haplotype diversity (**Table 3**; **Table S9**). Substantial differences in locus size between haplotypes were found in 07744-094 (*PdR1c*^+^/*PdR1*^-^) (1,103.9 and 842.8 kbp for Haplotype 1 and 2, respectively) and b42-26 (*PdR1*^-^/*PdR1*^-^) (969.7 and 848.1 kbp on Haplotype 1 and 2, respectively). In contrast, the two *PdR1* haplotypes of b43-17 (*PdR1a*^+^*/ PdR1b*^+^) differed by only 1.7 kbp (889.4 and 887.7 kbp on Haplotype 1 and 2, respectively), and the two haplotypes of b46-43 (*PdR1e*^+^*/PdR1f*^+^) were identical except for a single SNP located in a repetitive region at 472,149 bp after VMCNg2b7.2. Because our aim is to investigate the sequence diversity among *PdR1* haplotypes, only the Haplotype 1 of b46-43 was retained for the subsequent analyses, and named *PdR1e*.

**Table 3:**
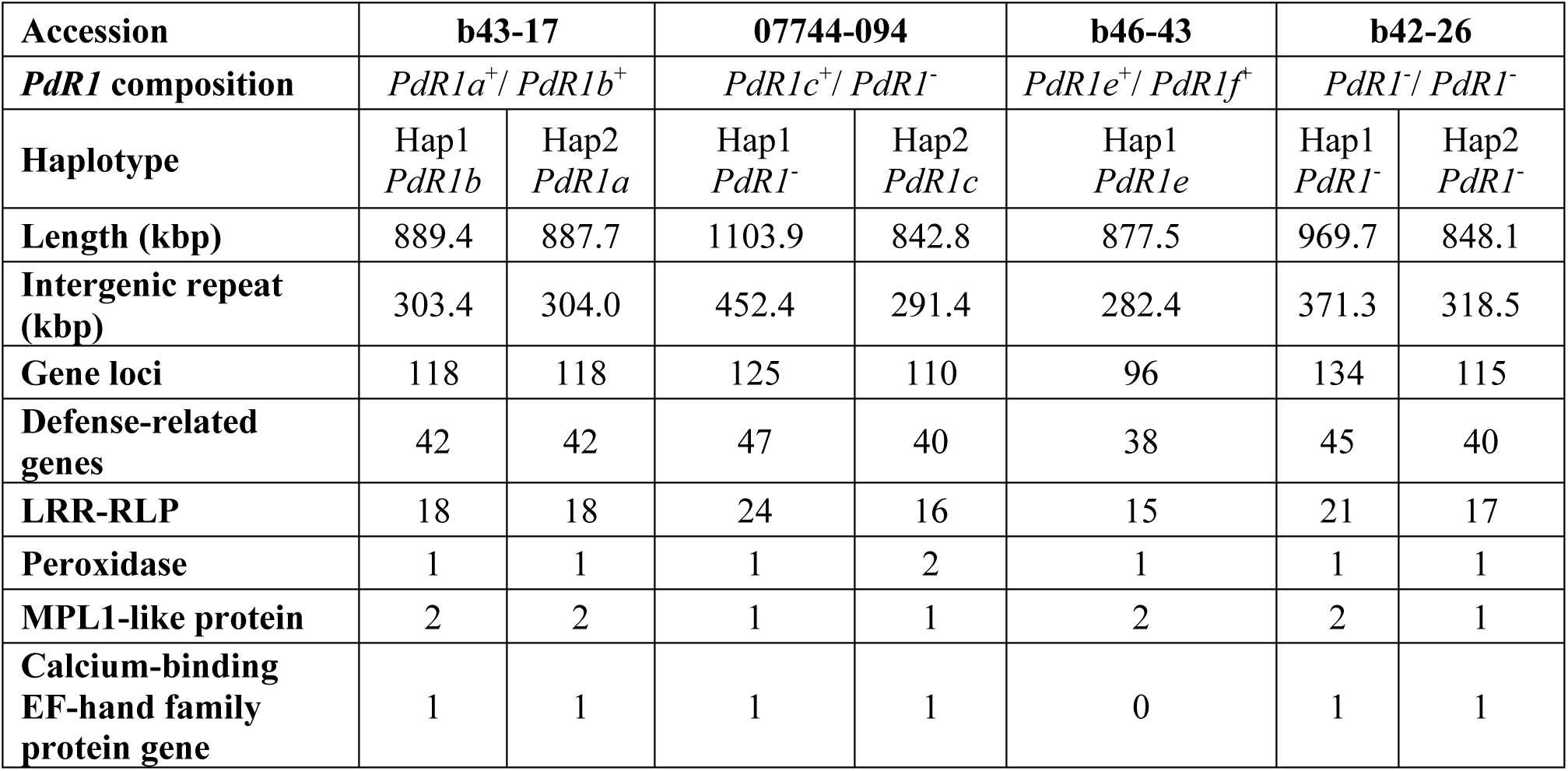
Features of the *PdR1* and *PdR1*^-^ haplotypes. Defense-related genes with different numbers across the haplotypes are depicted in this table. Number of defense-related genes composing each haplotype can be retrieved in Table S11. Gene content of each haplotype and their defense-related category and subcategory is detailed in Tables S12-S18. Abbreviations: Hap, Haplotype; LRR-RLP, leucine-rich repeat receptor-like proteins; MPL1, *Myzus persicae*-induced lipase 1.

To distinguish *PdR1c* in the backcross line 07744-094 (*PdR1c*^+^/*PdR1*^-^) from its homologous *V. vinifera PdR1*^-^ haplotype, we evaluated the normalized median base coverage (per 10 kbp) of b40-14 (*PdR1c*^+^/*PdR1d*^+^) on both haplotypes using short DNA-seq reads. Because 07744-094 is a backcross to *V. vinifera*, one haplotype derives from *V. arizonica* (carrying *PdR1c*) and the other from *V. vinifera* (*PdR1*^-^). Reads of b40-14 covered the entire region at 1.1 ± 0.3x on the Haplotype 2 of chromosome 14, whereas Haplotype 1 showed little to no coverage (**Fig. 1A**). This indicates that *PdR1c* is located on Haplotype 2 of 07744-094 genome, while the Haplotype 1 corresponds to the *V. vinifera PdR1*^-^ haplotype. Interestingly, coverage on Haplotype 2 between markers Ch14-77 and Ch14-02 (26.8 - 27.01 Mbp) averaged 1.5 ± 0.2x, suggesting that this region is highly homozygous between *PdR1c* and *PdR1d*.

**Fig. 1:**
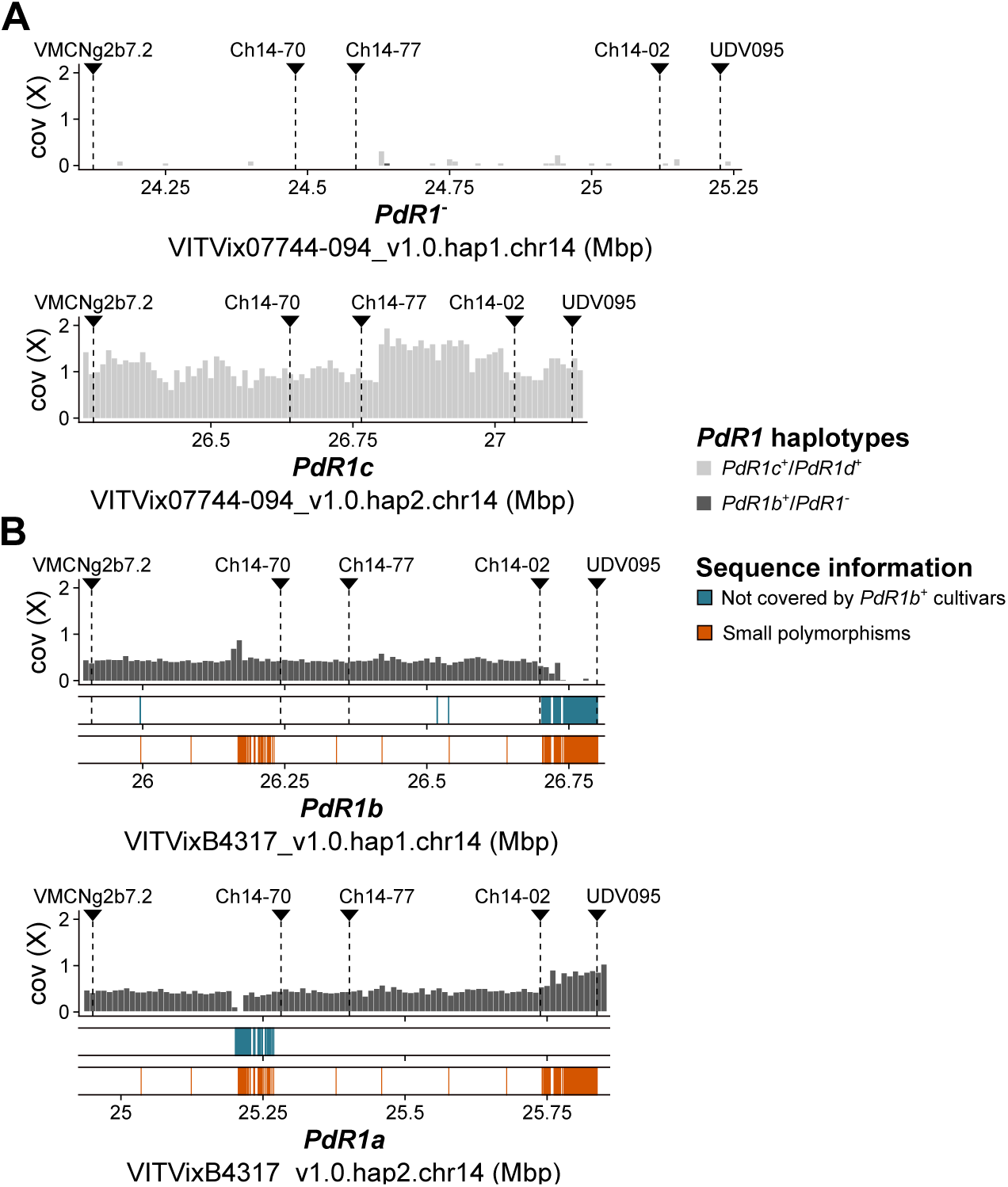
*PdR1* haplotyping of 07744-094 (*PdR1c*^+^/*PdR1*^-^) and b43-17 (*PdR1a*^+^*/ PdR1b*^+^). (**A**) Normalized median DNA-seq coverage per 10 kbp of b40-14 (*PdR1c*^+^/*PdR1d*^+^) on the diploid genome of 07744-094 (*PdR1c*^+^/*PdR1*^-^). (**B**) Average of the normalized median DNA-seq coverage per 10 kbp of five *PdR1b*^+^ cultivars on the genome of b43-17 (*PdR1a*^+^*/ PdR1b*^+^). Bases not covered by any *PdR1b*^+^ cultivar, and small polymorphisms between the two haplotypes of b43-17 (*PdR1a*^+^*/ PdR1b*^+^) are depicted for each haplotype. Chromosomal position of the *PdR1*-associated genetic markers is indicated by black triangles and dashed lines.

Similarly, we assessed the normalized median base coverage of five PD-resistant BC_4_ and BC_5_ cultivars carrying only *PdR1b* (Morales-Cruz et al., 2023) to distinguish *PdR1b* from *PdR1a* in b43-17 (**Fig. 1B**; **Figure S1-S2**). The average coverage across the five *PdR1b*^+^ cultivars was 0.4 ± 0.1x between markers VMCNg2b7.2 and Ch14-02 on both haplotypes of the b43-17 genome, suggesting that the region is highly homozygous in *PdR1a* and *PdR1b*. In the 3’ portion of the locus (between the markers Ch14-02 and UDV095), the Haplotype 1 showed little to no coverage, whereas Haplotype 2 showed 0.7 ± 0.1x coverage, suggesting that the five PD-resistant cultivars possess this part of the locus. In addition, a short region between markers VMCNg2b7.2 and Ch14-70 (from 25.2 to 25.21 Mbp) was uncovered in Haplotype 2, while a coverage peak (∼0.8x) was observed at 26.17 Mbp on the same haplotype. The localized differences in coverage between haplotypes could reflect either recombination or assembly-related haplotype switches. Remarkably, small polymorphisms (< 50 bp) between the two haplotypes of b43-17 were mostly located in the same two regions (**Fig. 1B**). Furthermore, no sign of haplotype switch in these two regions could be observed on the alignment of the HiFi reads of b43-17 on the genome (**Figures S4-S5**). Altogether, these results indicate that a recombination event occurred in the 3’ of the locus during the breeding of the five PD-resistant cultivars, and that *PdR1a* and *PdR1b* are located on the Haplotype 2 and Haplotype 1 of the chromosome 14 of b43-17, respectively.

### *PdR1* repeat and gene content

To determine the origin of *PdR1* haplotype size variation, we annotated the repeat and gene content of the four genomes. Annotation of repetitive elements showed that the total length of intergenic repeats in *PdR1* haplotypes ranged from 452.4 kb in the *V. vinifera PdR1*^-^ to 303.4 kb in *PdR1b* of b43-17 (**Table 3**, **Table S10**). These data indicate that *PdR1* haplotype size differences are primarily driven by variation in intergenic repeat content.

The number of annotated gene loci ranged from 96 in *PdR1e* to 134 in the *PdR1*^-^ of b42-26 Haplotype 1. Based on protein domain composition and homology to *A. thaliana* proteins, we identified genes potentially involved in plant responses to biotic stress in each haplotype (**Tables S11–S18**). Defense-related genes within *PdR1* haplotypes included those encoding extracellular receptors and transcription factors, as well as genes involved in reactive oxygen species (ROS) production and scavenging, cell wall modification, programmed cell death, ethylene- and jasmonic acid–mediated signaling, and genes with putative roles in biotic stress responses of unknown mechanism (**Table S11**). The number of defense-related genes ranged from 38 in *PdR1e* to 47 in the *V. vinifera PdR1*^-^ (**Table 3**; **Table S11**).

Differences in defense-related gene content among *PdR1* haplotypes were mainly driven by variation in genes encoding LRR-RLPs between markers Ch14-77 and Ch14-02, which ranged from 16 to 24 genes. In addition, *PdR1c* contained two peroxidase-coding genes, compared with a single gene in the other haplotypes. Two genes encoding homologs of *Myzus persicae*–induced lipase 1 (MPL1) from *A. thaliana* were present in *PdR1a*, *PdR1b*, *PdR1e*, and the *PdR1*^-^ Haplotype 1 of b42-26, whereas only one *MPL1* homolog was detected in the remaining haplotypes. In *A. thaliana*, MPL1 restricts *Fusarium graminearum* infection by limiting jasmonic acid accumulation (Alam et al., 2022). A gene encoding a calcium-binding EF-hand family protein was present in all haplotypes except *PdR1e*.

Altogether, variation in intergenic repeat and gene composition, particularly among defense-related genes, provides clear evidence of sequence diversity between *PdR1* and *PdR1^-^* haplotypes, as well as among the *PdR1* haplotypes themselves.

### *PdR1* sequence graph

To further evaluate sequence diversity among the *PdR1* haplotypes, we constructed a sequence graph including both *PdR1* and *PdR1^-^* haplotypes. This graph comprised 92,807 nodes spanning 1,442,276 bp. An extensive and complex bubble located approximately three-quarters along the graph indicated divergent paths consistent with large structural variation in this region (**Fig. 2A**). Several smaller bubbles elsewhere in the graph reflected finer-scale polymorphisms, indicating substantial structural divergence concentrated in a specific region of the locus, together with additional localized variation among haplotypes.

**Fig. 2:**
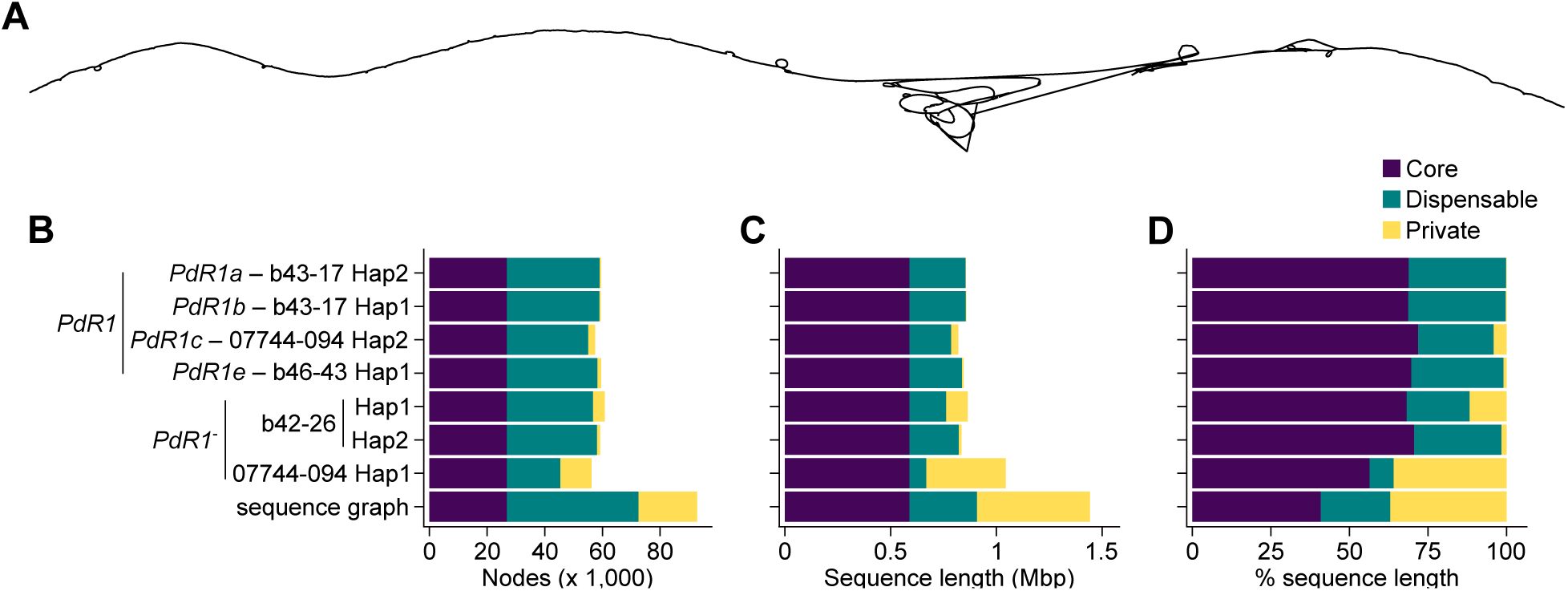
Sequence graph of the *PdR1* and *PdR1*^-^ haplotypes. (**A**) 2D visualization of the sequence graph built using 4 *PdR1* haplotypes: *PdR1a* and *PdR1b* of b43-17, *PdR1c* from 07744-094 (*PdR1c*^+^/*PdR1*^-^), *PdR1e* of b46-43, and 3 *PdR1*^-^ haplotypes: the two haplotypes of b42-26 (*PdR1*^-^/*PdR1*^-^), and the *V. vinifera* haplotype of 07744-094. Number of nodes (**B**), cumulative sequence length (**C**), and percentage of sequence length (**D**), made of core, dispensable, and private nodes in each haplotype path. Panels **B–D** share the same legend. Abbreviations: Hap, Haplotype.

To characterize this variation, we classified graph nodes into three categories: core nodes (shared across all seven haplotypes), dispensable nodes (present in two to six haplotypes), and private nodes (specific to a single haplotype). Nearly half of the nodes were dispensable (49.1%), whereas core and private nodes accounted for 29.0% and 21.9%, respectively (**Fig. 2B**). Private nodes were most abundant in the *V. vinifera PdR1*^-^ haplotype (10,711 nodes), and, after core nodes, represented the largest contribution to node content, followed by nodes shared among *V. arizonica* and its hybrids (**Fig. S4**). This pattern indicates marked sequence divergence between *V. vinifera* and *V. arizonica*–derived haplotypes in this region. Among *PdR1* haplotypes, *PdR1c* contained the greatest number of private nodes (2,334), followed by *PdR1e* (1,253), and the two haplotypes of b43-17 (476 and 550). Although fewer than dispensable nodes, private nodes accounted for a greater proportion of total graph length (37% *vs*. 22.2%) (**Fig. 2C & D**), indicating that private nodes tend to be longer than shared nodes.

Comparison of *PdR1* and *PdR1^-^* haplotype paths identified 8,562 *PdR1*-specific nodes, defined as nodes present in at least one *PdR1* haplotype and absent from all *PdR1^-^* haplotypes, representing 9.2% of all graph nodes. All four *PdR1* haplotype paths contained *PdR1*-specific nodes (**Fig. 3A**), ranging from 3,139 in *PdR1e* to 4,295 in *PdR1b*. The two b43-17 haplotypes contained the highest numbers of *PdR1*-specific nodes (3,479 and 3,505 in *PdR1a* and *PdR1b*, respectively). Most *PdR1*-specific nodes were shared among multiple haplotype paths, particularly between the two b43-17 haplotypes and *PdR1e*. In contrast, *PdR1c* exhibited a distinct pattern, with 71.8% of its *PdR1*-specific nodes being private, accounting for the largest cumulative private-node length (33.1 kbp) (**Fig. 3B**).

**Fig. 3:**
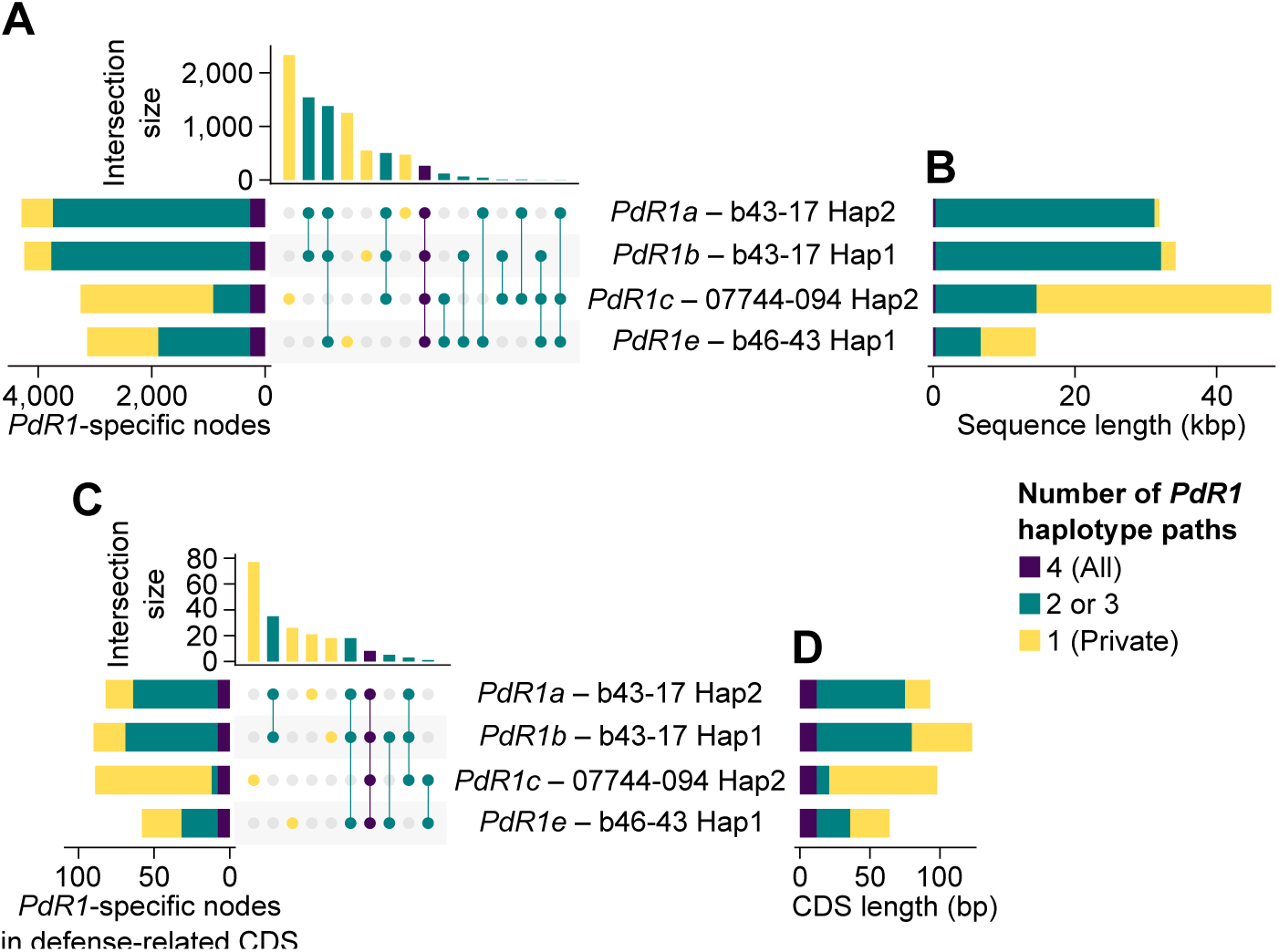
Nodes specific to the paths of the *PdR1* haplotypes in the sequence graph. (**A**) Number of *PdR1*-specific nodes (*i.e*. found only in the paths of the *PdR1* haplotypes) in each *PdR1* haplotype path, and their distribution among the paths. (**B**) Cumulative sequence length of the *PdR1*-specific nodes in each haplotype path. Number of *PdR1*-specific nodes overlapping the protein-coding sequence (CDS) of the genes potentially involved in biotic stress response, their repartition among the paths (**C**), and their cumulative length (**D**). Panels **A–D** share the same legend. Abbreviations: Hap, Haplotype.

Among *PdR1*-specific nodes, 212 overlapped protein-coding sequences (CDSs) of defense-related genes, corresponding to 2.5% of all *PdR1*-specific nodes. These nodes were detected across all four *PdR1* haplotype paths, with the highest numbers in *PdR1b* and *PdR1c* (90 and 89, respectively), followed by *PdR1a* (82) and *PdR1e* (58) (**Fig. 3C**). Overall, these nodes overlapped approximately half of the defense-related gene content of each *PdR1* haplotype, ranging from 52.4% to 53.7%. They affected all defense-related gene categories, albeit in different proportions among haplotype paths (**Table S19**). Only eight nodes were shared across all four *PdR1* haplotypes, six of which overlapped genes involved in cell wall modification (**Fig. 3C**, **Table S19**).

Most *PdR1*-specific, defense-related nodes were private to *PdR1c* (86.5%), whereas nodes in the two b43-17 haplotypes were largely shared between each other and with *PdR1e*. In *PdR1c*, private nodes predominantly overlapped CDSs of LRR-RLP genes (45 nodes) and genes involved in ROS production (16 nodes) (**Table S19**). *PdR1a* and *PdR1b* shared nodes overlapping genes encoding extracellular receptors (LRR-RLPs (14) and LRR-RLK (2)), the metal–nicotianamine transporter YSL6-like protein (9), and cell wall modification genes (7). In contrast, *PdR1e* shared 24 *PdR1*-specific, defense-related nodes with one or two other *PdR1* haplotypes, mostly with the b43-17 haplotypes, and contained 26 private nodes, which overlapped genes associated with cell death and genes with putative roles in biotic stress responses of unknown mechanism.

In summary, the sequence graph analysis revealed that each *PdR1* haplotype path contains *PdR1*-specific nodes, but with distinct distributions, lengths, and functional enrichments, affecting different categories of defense-related genes.

### *PdR1c* candidate genes

Because the assembly of 07744-094 (*PdR1c⁺/PdR1⁻*) enabled unambiguous phasing of *PdR1c* and the sequence graph revealed *PdR1*–specific features of the region, we investigated the *PdR1c* gene content to identify candidate genes associated with PD resistance. We previously refined the boundaries of the *PdR1* region in *V. arizonica* b40-14 between the markers Ch14-77 and Ch14-02 (Morales-Cruz et al., 2023). In *PdR1c*, these markers delimited a region spanning 26,764,799–27,034,556 bp on chromosome 14 of Haplotype 2 of 07744-094. This refined region encompassed 39 genes, including 18 defense-related genes encoding a chloride channel protein, 16 LRR-RLPs, and an MPL1-like protein (**Fig. 4**, **Table S14**). Sequence graph analysis identified 709 *PdR1*-specific nodes within this interval, 97% of which were private to *PdR1c* (**Table S20**).

**Fig. 4:**
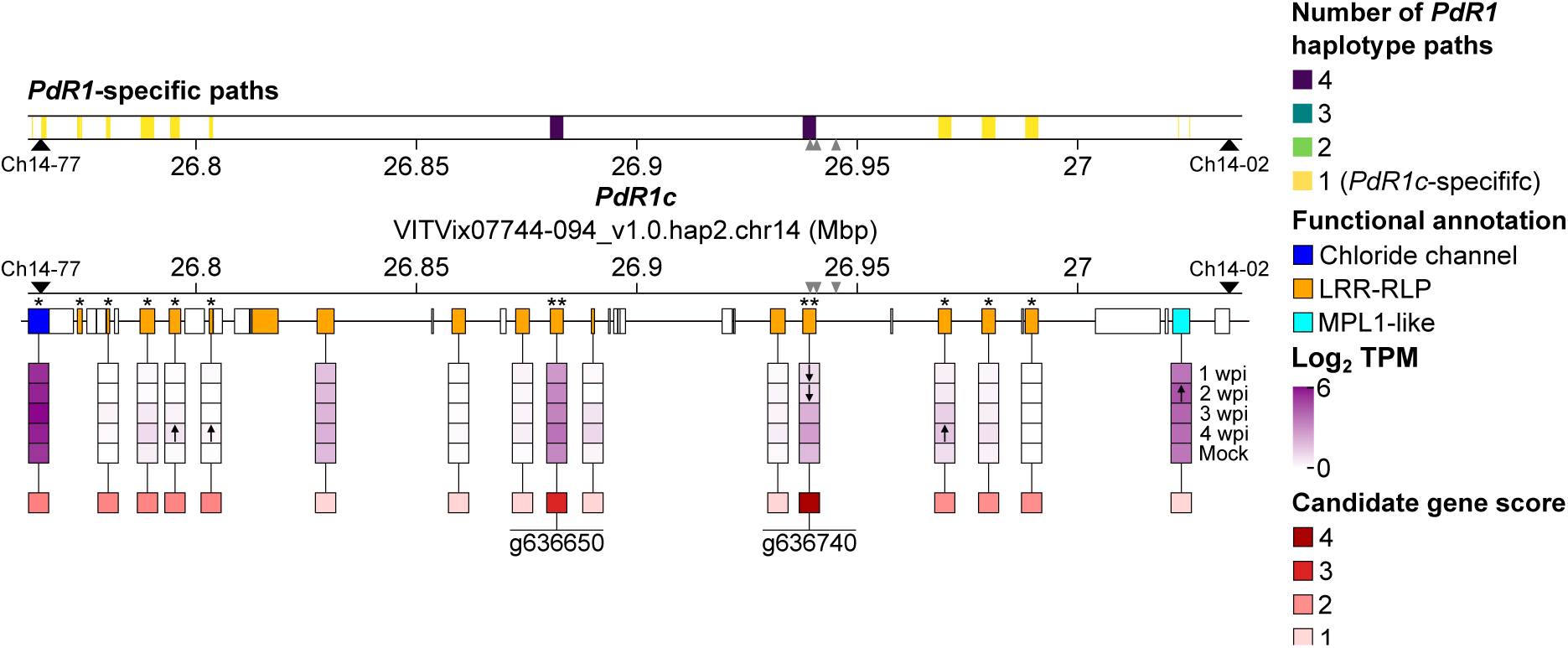
Narrowing candidate *PdR1c* genes using sequence graph information and gene expression. *PdR1*-specific paths in the defense-related genes of the refined *PdR1c* region. Gene content of the *PdR1c* locus with the defense-related genes colored according to their functional annotation. Genes with a protein sequence specific to *PdR1c* are indicated with *, genes with a protein sequence specific to all *PdR1* haplotypes are depicted with **. Expression of the defense-related genes in stems of 3 PD-resistant 07744 lines at 1, 2, 3, 4 weeks post *Xf* inoculation (wpi) and stems mock-inoculated (Mock). Gene expression is represented as the log_2_-transformed mean Transcripts per Million (TPM) for the genes expressed in at least one condition (TPM > 0). Genes more highly and lowly expressed are indicated by ↑ and ↓, respectively. Score of the defense-related genes of the refined *PdR1c* region as candidate genes. Scoring was made based on the position of the genes regarding the PD resistance–associated kmers, the specificity of their protein sequence to all *PdR1* haplotypes and *PdR1c*, and gene expression. Black triangles represent the position of the *PdR1*-associated genetic markers. Gray triangles indicate the positions of the PD resistance–associated kmers.

To assess the functional impact of *PdR1*-specific variation, we compared CDS paths of defense-related genes in *PdR1c* with those of the three other *PdR1* and three *PdR1^-^* haplotypes (**Fig. 4**). *PdR1*-specific CDS paths were detected in 12 defense-related genes, encoding the chloride channel protein, 10 LRR-RLPs, and the MPL1-like protein. All *PdR1*-specific CDS paths were unique to *PdR1c*, except for two monoexonic LRR-RLP genes (g636650 and g636740) (**Table S21**). Comparison of CDS paths and predicted protein sequences showed that nine genes encoded *PdR1c*-specific proteins, including the chloride channel protein gene and eight LRR-RLP genes, whereas g636650 and g636740 encoded identical protein sequences across all *PdR1* haplotypes. Notably, g636740 overlapped PD resistance–associated kmers, previously identified using 167 *V. arizonica* accessions and hybrids, providing independent support for its association with PD resistance (Morales-Cruz et al., 2023). Forty-six kmers aligned perfectly (**Table S22**), spanning 26,939,281–26,945,206 bp within the Ch14-77–Ch14-02 interval, which contains only the CDS of g636740.

To further narrow down candidate genes, we profiled the expression of defense-related genes in *Xf-*inoculated stems from three PD-resistant 07744 backcross lines, including 07744-094, at 1, 2, 3, and 4 weeks post inoculation, as well as in mock-inoculated controls. Sixteen of the 18 defense-related genes were expressed (TPM > 0) in at least one condition (**Fig. 4**). These included the chloride channel protein, 14 LRR-RLPs, and the MPL1-like gene. Only four genes showed differential expression in response to *Xf*, comprising three LRR-RLP genes and the MPL1-like gene.

Based on these results, we scored defense-related genes within the refined *PdR1c* region as candidate PD resistance genes using three criteria: (i) proximity to PD resistance–associated kmers, (ii) protein sequence specificity to all *PdR1* haplotypes and/or *PdR1c*, and (iii) constitutive expression and/or transcriptional response to *Xf* infection (**Fig. 4; Table S23**). The LRR-RLP gene g636740 had the highest score (4), followed by the LRR-RLP gene g636650 (3). Eight genes encoding the chloride channel protein and 7 LRR-RLPs had a score of 2, while 6 genes coding for the MPL1-like and 5 LRR-RLPs had a score of 1. Accordingly, LRR-RLP genes g636740 and g636650 are the strongest *PdR1c* candidates.

## Discussion

To evaluate the sequence diversity among the *PdR1* haplotypes from *V. arizonica* and its hybrids, we first assembled and haplotyped the diploid genome of four PD-resistant accessions: 3 with their PD resistance associated with *PdR1* (b43-17 (*PdR1a*^+^*/ PdR1b*^+^), the backcross 07744-094 (*PdR1c*^+^/*PdR1*^-^), and b46-43 (*PdR1e*^+^*/PdR1f*^+^)), and b42-26 (*PdR1*^-^/*PdR1*^-^) which displays quantitative PD resistance but not associated with *PdR1* (**Table 1**) (Riaz et al., 2018). Genome assembly using HiFi reads allowed to generate all *PdR1* and *PdR1*^-^ haplotypes in a single contig, which is an improvement for *PdR1c* compared to two fragmented loci in the genome assembly of *V. arizonica* b40-14 (*PdR1c*^+^*/PdR1d*^+^) (**Figure S1**). Localization of the *PdR1* region through the *in silico* amplification of *PdR1*-associated genetic markers showed differences in amplicon size and locus length between accessions and within each accession, except in b46-43 which both haplotypes identical at the exception of 1 SNP. Differences were also found in the gene and repeat contents of the *PdR1* haplotypes, especially in the number of LRR-RLP genes. All these differences provide clear evidence of sequence diversity between *PdR1* and *PdR1*^-^ haplotypes, as well as among the *PdR1* haplotypes themselves.

Sequence diversity was further investigated through the construction of a sequence graph including 7 haplotypes. These consisted in 4 *PdR1* haplotypes: *PdR1a*, *PdR1b*, *PdR1c*, and *PdR1e*, and 3 *PdR1*^-^ haplotypes: one from *V. vinifera* and two from the PD-resistant accession b42-26. The 2D representation of the sequence graph (**Fig. 2A**) showed an extensive structural divergence located approximately three-quarters along the graph, together with additional localized structural variations among haplotypes. Categorization of the graph nodes as core, dispensable, and private, determined a substantial sequence divergence between *V. vinifera* and *V. arizonica*–derived haplotypes in the locus (**Fig. 2B–D**). Interspecific sequence diversity has also been observed in other grape disease resistance loci, such as the powdery mildew-resistance loci *Run1.2* and *Run2.2* (Cochetel et al., 2021), *Ren6* and *Ren7* (Massonnet et al., 2025), and the downy mildew-resistance loci *Rpv3* (Wilkerson et al., 2025). Using the inter- and intra-specific sequence diversity between the 4 *PdR1* haplotypes and the *V. vinifera PdR1*^-^ haplotype of 07744-094 and the two haplotypes of b42-26, we determined the nodes specific to the *PdR1* haplotypes (*i.e*. found in at least one *PdR1* haplotype but absent in all *PdR1*^-^ haplotypes). Investigating the *PdR1*-specific nodes showed a diversity in sequence between the four *PdR1* haplotypes (**Fig. 3**), while examining their overlap with defense-related genes showed that they affected different defense-related gene categories.

Finally, we demonstrated the utility of the sequence graph to narrow down candidate genes in the refined region of *PdR1c*, especially when coupled with gene expression. Scoring of the expressed defense-related genes within the refined *PdR1c* region based on proximity to PD resistance–associated kmers, protein sequence specificity, and gene expression, pointed two strong *PdR1c* candidates: the LRR-RLP genes g636740 and g636650 (**Fig. 4**). The same approach could be used for the 3 other *PdR1* haplotypes through the profiling of the gene expression of their defense-related genes in petioles or stems at both constitutive level and in response to *Xf*. This could help to determine whether the expression of the LRR-RLP genes g636740 and g636650 is common in *PdR1*^+^ accessions. Two LRR-RLP genes were also proposed as candidate genes in *V. arizonica* b40-14 (Morales-Cruz et al., 2023). These genes correspond to g636600 and g636650 in 07744-094 genome. Based on the results of our sequence graph, g636650 is a very strong candidate as its protein sequence is identical for all the *PdR1* haplotypes and absent in all *PdR1*^-^ haplotypes. However, the protein sequence of g636600 was also found present in b42-26, decreasing its score as candidate gene. Gene expression profiling of *Xf*-infected tissues of b42-26 could help to determine whether g636600 is still a strong candidate. Regarding g636740, the gene was absent in the genome of *V. arizonica* b40-14. In b40-14 genome, the refined *PdR1* region was made of 2 and 4 contigs on Haplotype 1 and 2, respectively (Morales-Cruz et al., 2023)(**Figure S1**). The fragmentation of the locus region and the absence of haplotyping information for the genome assembly are likely at the origin of the absence of g636740 in the genome of *V. arizonica* b40-14.

Future work will focus on the functional validation of the *PdR1c* candidate genes, the incorporation of additional *PdR1* haplotypes in the sequence graph, and the profiling of the gene expression of the defense-related genes in several *PdR1* haplotypes, at both constitutive level and in response to *Xf*. This would help to determine whether the PD resistance associated with the four *PdR1* haplotypes is based on the expression of the same gene(s). As a protocol for stable protoplast-based gene editing has been developed for *V. arizonica* (Tricoli and Debernardi, 2024), stable gene editing of the *PdR1c* candidate genes in *V. arizonica* b40-14 (*PdR1c*^+^*/PdR1d*^+^) could be an option. However, the lack of sequence information about *PdR1d* might complicate the process. Stable expression of the candidate genes in *V. vinifera* could be another option to assess the role of these genes in PD resistance. On the other hand, using transient transformation methods, such *as Agrobacterium*-mediated expression (Zhang et al., 2024), virus-induced gene silencing (Yang et al., 2022), and exogenous dsRNA-based RNAi (Marcianò et al., 2021), would likely be more challenging to evaluate the function of the candidate genes as *Xf* is a xylem-restricted pathogen.

From a breeding perspective, the comparison of the *PdR1* and *PdR1*^-^ haplotypes through the sequence graph could greatly help the development of precise markers either specific to all *PdR1* haplotypes or specific to each single haplotype. Such markers could accelerate the screening in traditional breeding programs or the incorporation of the genes responsible for PD resistance in elite cultivars using gene editing (Greenwood et al., 2023).

## Data availability

HiFi reads are accessible through NCBI under the BioProject PRJNA1380937. Genome sequences and their annotation files are available at Zenodo doi: 10.5281/zenodo.17925270. These files will be released upon acceptation of the manuscript. HiFi reads can be accessed using the following reviewer link: https://dataview.ncbi.nlm.nih.gov/object/PRJNA1380937?reviewer=8irh5k96c575be0sufv7de61 <u>sg</u>. Access to genome and annotation files can be requested at Zenodo doi: 10.5281/zenodo.17925270.

## Acknowledgments

We thank the UC Davis DNA Technologies Core for sequencing assistance.

## Author contributions

Conceptualization: D.C. Formal analysis: M.M., M.Z, N.C. Investigation: M.M., M.Z, R.F.-B, N.C. Resources: S.R. Writing—original draft preparation: M.M., D.C. Writing—review and editing: M.M., D.C. Visualization: M.M. Project administration: D.C. Funding acquisition: D.C, S.R. All authors read and agreed to the published version of the manuscript.

## Supplementary Information

### Tables

**Table S1**: Genome sequencing summary.

**Table S2**: Summary of the draft genome assemblies.

**Table S3**: Summary of the draft genome assemblies.

**Table S4**: Functional annotation of the predicted proteins of b43-17 (*PdR1a*^+^*/PdR1b*^+^).

**Table S5**: Functional annotation of the predicted proteins of 07744-094 (*PdR1c*^+^*/PdR1*^-^).

**Table S6**: Functional annotation of the predicted proteins of b46-43 (*PdR1e*^+^*/PdR1f*^+^)

**Table S7**: Functional annotation of the predicted proteins of b42-26 (*PdR1*^-^*/PdR1*^-^)

**Table S8**: Coordinates of the *PdR1*-associated marker amplicons in the studied genomes.

**Table S9**: Chromosomal position of the *PdR1* and *PdR1*^-^ haplotypes.

**Table S10**: Intergenic repeat content of the *PdR1* and *PdR1*^-^ haplotypes.

**Table S11**: Defense-related gene content of the *PdR1* and *PdR1*^-^ haplotypes.

**Table S12**: Functional annotation of the predicted proteins composing *PdR1a* of *b43-17* (*PdR1a*^+^*/PdR1b*^+^).

**Table S13**: Functional annotation of the predicted proteins composing *PdR1b b43-17* (*PdR1a*^+^*/PdR1b*^+^).

**Table S14**: Functional annotation of the predicted proteins composing *PdR1c* of 07744-094 (*PdR1c*^+^*/PdR1*^-^).

**Table S15**: Functional annotation of the predicted proteins composing *PdR1e* of *b46-43* (*PdR1e*^+^*/PdR1f*^+^).

**Table S16**: Functional annotation of the predicted proteins composing the *PdR1*^-^ haplotype of 07744-094 (*PdR1c*^+^*/PdR1*^-^).

**Table S17**: Functional annotation of the predicted proteins composing the *PdR1*^-^ haplotype of Haplotype 1 of b42-26 (*PdR1*^-^*/PdR1*^-^).

**Table S18**: Functional annotation of the predicted proteins composing the *PdR1*^-^ haplotype of Haplotype 2 of b42-26 (*PdR1*^-^*/PdR1*^-^).

**Table S19**: Number of *PdR1*-specific nodes overlapping the coding sequence of the defense-related genes made of nodes present in all four paths, 2 or 3 paths, a unique path (private), in each *PdR1* haplotype path.

**Table S20**: Chromosomal position of the *PdR1c*-specific nodes in the refined region.

**Table S21**: Comparison of the paths of defense-related CDS composing the *PdR1c* with the other haplotype paths, and the alleles of each gene. Paths that contained the same path than *PdR1c* are represented with a 1, while 0 indicates the absence of the paths.

**Table S22**: Chromosomal position of the *V. arizonica* PD resistance-associated kmers in *PdR1c*.

**Table S23**: Score of the defense-related genes of the refined *PdR1c* as candidate genes based on the position of the genes regarding the PD resistance–associated kmers, the specificity of their protein sequence to all *PdR1* haplotypes and *PdR1c*, and gene expression.

### Figures

**Figure S1: Contig composition of the two *PdR1* haplotypes in *V. arizonica* b40-14.** Contigs composing each *PdR1* haplotype in the genome of *V. arizonica* b40-14 (VITVarB40-14_v2.0.fasta).

**Figure S2: Normalized median base coverage per 10 kbp of five backcross cultivars carrying *PdR1b* at the *PdR1* locus of b43-17 Haplotype 1.** Only DNA-seq reads out of repetitive elements and aligning perfectly were used for base coverage analysis.

**Figure S3: Normalized median base coverage per 10 kbp of five backcross cultivars carrying *PdR1b* at the *PdR1* locus of b43-17 Haplotype 2.** Only DNA-seq reads out of repetitive elements and aligning perfectly were used for base coverage analysis.

**Figure S4: Alignment of the HiFi reads of b43-17 at the two heterozygous regions in the *PdR1* region of the Haplotype 1.** Only INDELs with a size greater that 100 bp are represented.

**Figure S5: Alignment of the HiFi reads of b43-17 at the two heterozygous regions in the *PdR1* region of the Haplotype 2.** Only INDELs with a size greater that 100 bp are represented.

**Figure S6: Comparison of the nodes composing the path of each haplotype in the *PdR1* sequence graph.**

